# Use of extracellular polymer substances as additives for improving biogas yield and digestion performance

**DOI:** 10.1101/699314

**Authors:** Haitong Ma, Chen Yan Guo, Ming Wu, Hui Liu, Zhiwei Wang, Shuangfei Wang

**Affiliations:** College of Light Industry and Food Engineering, Guangxi University, Nanning 530004, PR China; Key Laboratory of Clean Pulp and Paper and Pollution Control, Nanning 530004, PR China; College of Environmental Science and Forestry, State University of New York, Syracuse, NY 13210, USA

**Keywords:** extracellular polymer substances, methanogenesis, anaerobic digestion, biomass conductive materials

## Abstract

To understand how extracellular polymer substances (EPS) as additives promotes methanogenesis, batch tests of methane production potential in anaerobic reactors with the addition of EPS or not were conducted. Research showed that EPS increased remarkably methane production during anaerobic digestion (36.5% increase compared with the control). EPS enriched functional microorganisms such as *Firmicutes, Actinobacteria, Synergistetes*, and *Chloroflexi*. Among them, 8.86% OTUs from the important hydrolysis and acidification phyla, which may be an important reason for the enhanced methanogenic capacity of anaerobic granular sludge. Additionally, EPS also improved the abundance of cytochrome c (c-Cyts), accelerating the direct interspecies electron transfer (DIET) between syntrophic bacteria and methanogens, thus enhancing the methane production. Interestingly, the average particle size, volatile suspended solids/total suspended solids (VSS/TSS) and EPS content of anaerobic granular sludge (AnGS) in the EPS reactor were approximately equal to that of the control reactor during the anaerobic digestion, illustrating that EPS could not affect the physicochemical properties of AnGS. Therefore, these results suggested that EPS mainly played a role in the form of conductive materials in the anaerobic digestion process. Compared with conductive materials, EPS as biomass conductive materials was not only environmentally friendly and economical but also no secondary pollution.

**Importance:** Compared with the reported conductive materials, EPS has the potential of biodegradation, electron transfer and no significant secondary pollution. Besides, there are few studies on the utilization of EPS resources, especially the effect of EPS as an additive on anaerobic digestion performance. To clarify whether EPS as conductive materials or carbon source promotes methanogenesis. Therefore, in this study, we investigated the influence of EPS as an additive on the methanogenic capacity, physical-chemical properties, microbial community structure and metabolic function of anaerobic granular sludge (AnGS), and preliminarily expatiate the influence mechanism of EPS as an additive on methanogenesis. At the meantime, the research is expected to provide new solutions for the improvement of anaerobic digestion performance and disposal of waste mud.

## Introduction

Anaerobic digestion (AD) is a significant process that converts organic waste into clean energy (CH_4_) by utilizing the metabolism of microorganisms, which has attracted extensive attention because of its potential to generate bioenergy (Batstone & Virdis, 2014; Liu et al., 2016). However, the low efficiency, time-consuming, and low methane yield limit its application in the industrial wastewater treatment (Zhang et al., 2017).

At present, using conductive materials (carbon-based materials and conductive minerals) to improve the anaerobic digestion performance is a research hotspot. For example, nano-zero-valent iron (Suanon et al., 2017), goethite (Tan et al., 2015), hematite (Kato et al., 2012) and magnetite (Zhuang et al., 2015) can significantly increase methane production in anaerobic digesters by facilitating direct electron transfer between fermenting bacteria and methanogens or enriching functional microorganisms. Direct interspecies electron transfer (DIET) is an unconventional syntrophic methanogenesis metabolism in which electrons flow from syntrophic bacteria to methanogens via cell components like cytochrome c (c-Cyts), pilin, etc (Dubé & Guiot, 2015; Lovley, 2016). Generally, while considering the diffusion limitation of the substrate, the DIET performs obviously faster than methanogenesis via hydrogen or formate during the anaerobic digestion (Cruz et al., 2014; Zhang et al., 2017). In addition, carbon-based materials like granular activated carbon (Liu et al., 2012; Zhang et al., 2017), graphite (Zhao et al., 2015) and biochar (Yu et al., 2015) were also reported to enhance the anaerobic digestion efficiency by DIET. It should be emphasized that these non-biodegradable conductive minerals and carbon-based materials are not only complicated in the production process, high in cost and difficult to recycle, but also easy to cause secondary pollution. More importantly, the addition of conductive materials could increase the difficulty of subsequent treatment.

Extracellular polymer substances (EPS) is a natural polymer substance secreted by microorganisms and mainly composed of protein, polysaccharide, humus and DNA, distributing in the interior and surface of the sludge (Sheng et al., 2010). The reviewed literature found that EPS has an important influence on the sedimentation, dewatering, and flocculation of sludge, but due to the complexity of EPS and the lack of basic knowledge, its impact on the biochemical performance of the sludge remains to be further explored (Shi et al., 2017). Recently, Xiao et al. (2017) reported that EPS can store electrochemically active substances such as c-Cyts, humus and tightly-bound EPS, which stimulated DIET between fermenting bacteria and methanogens. Similarly, Ye et al. (2017) also found that the redox properties of EPS are closely related to the methanogenic capacity of the sludge in red mud reactors, and the higher the c-Cyts abundance, the faster the electron transfer rate between microorganisms. Furthermore, part of EPS can also be used by bacteria as sources of carbon and energy in conditions of nutrient shortage (Sutherland, 2001a; Zhang and Bishop, 2003). However, Wang et al. (2005, 2007) argued that certain parts of EPS cannot be degraded by microorganisms. These results provide the possibility for EPS to be used as additives to improve the anaerobic digestion performance. But it is still confused whether EPS as conductive materials or carbon source promotes methanogenesis.

Compared with the reported conductive materials, EPS has the potential of biodegradation, electron transfer and no significant secondary pollution. Besides, there are few studies on the utilization of EPS resources, especially the effect of EPS as an additive on anaerobic digestion performance. To clarify whether EPS as conductive materials or carbon source promotes methanogenesis. Therefore, in this study, we investigated the influence of EPS as an additive on the methanogenic capacity, physical-chemical properties, microbial community structure and metabolic function of anaerobic granular sludge (AnGS), and preliminarily expatiate the influence mechanism of EPS as an additive on methanogenesis. At the meantime, the research is expected to provide new solutions for the improvement of anaerobic digestion performance and disposal of waste mud.

## 2 Materials and Methods

### 2.1. Materials

EPS as an additive was extracted from the AnGS. The method was based on a chemical process in which a solution of formaldehyde (36.5%) and NaOH (1 mol/L) is added to the sludge, and the mixture is agitated at 200 rpm for 2 to 3 h, and then centrifuged at 10,000×g for 10 min, and eventually filtered through a 0.22 μm membrane to obtain the filtrate (Adav & Lee., 2008). Finally, the filtrate in the freeze drier freeze-dried, 4℃ saved for later use. AnGS was collected from a local paper mill (Nanning, China). In order to reduce the experimental error, AnGS with an average particle size of 0.3~0.6 mm was selected for seed sludge. The characteristics of the synthetic wastewater using glucose as carbon source: glucose (25.00 g/L), NaHCO_3_ (16.25 g/L), NH_4_Cl (3.00 g/L), KH_2_PO_4_ (0.56 g/L), FeSO_4_·4H_2_O (0.032 g/L), CaCl_2_ (0.038 g/L), MgSO_4_·7H_2_O (0.042 mg/L), and pH = 7.20 ± 0.20.

### 2.2 Anaerobic digestion experiment

A mass of 0.15 and 0.3 g of EPS was added into 250 mL anaerobic serum bottle to construct the EPS supplemented reactors (A and B) with 25 mL of AnGS and 150 mL of the substrate. The bottle without EPS addition was designated the control reactor (C). The bottles were emptied of air by the flush of nitrogen for 15 min and sealed with stoppers and tapes. Then, the reactors were maintained at (37 ± 1℃) with a constant temperature incubator for a period of 28 days. During the study, a simple drainage gas-collecting method was adopted to measure the daily biogas yield over 28 days. Before the biogas volume was measured, each reactor was shaken for 2 min to ensure the mixture was well-distributed, and the gas produced by reactors was passed through a 1.5% ~ 5% NaOH solution to remove acid gases. The samples for EPS, volatile suspended solids/total suspended solids (VSS/TSS) and average particle size analysis was obtained from anaerobic reactors every seven days. At the end of the experiment, the microbial community and function of AnGS were discussed by the high-throughput sequencing technology.

### 2.3 Analytical method

#### 2.3.1 VSS/TSS, biogas yield and average particle size

The average particle size of AnGS was analyzed by ultra-high-speed intelligent particle size analyzer (Mastersizer 3000). Analysis of VSS/TSS and biogas yield was conducted according to Standard Methods for the Examination of Water and Wastewater.

#### 2.3.2 Analysis of EPS

The method for EPS extraction was as described by Adav & Lee. (2008). The protein, humus and polysaccharide concentrations were determined by a modified Lowry method and the anthrone-sulfuric acid spectrophotometric method using bovine serum albumin, humic acid salt and glucose as the respective standards. UV/Visible (UV/Vis) spectroscopy in a diffuse transmission mode was used to measure the electronic absorption spectra of the multi-heme c-Cyts in the EPS (Ye et al., 2018). EEM fluorescence spectroscopy was used to characterize EPS in this research, and excitation and emission spectra were scanned from 220 to 550 nm in 5 nm increments and from 250 to 550 nm in 5 nm increments, respectively (Ding et al., 2015).

#### 2.3.3 DNA extraction, high-throughput sequencing and data analysis

To understand the change in the microbial communities in anaerobic reactors, the samples were collected and sent to Majorbio (Shanghai, China) for DNA extraction and amplicon sequencing according to Miseq protocols. The samples were ready for DNA isolation after the experiment ended, respectively tagged as C (collected from C reactor), B (collected from B reactor), and A (collected from A reactor). For high-throughput sequencing, the primer pairs (341F and 805R) were used as a universal gene primer to amplify the V4 region of the 16S rRNA gene with the barcode. The detailed sequencing process was as described by Zhang *et al.* (2017). Sequences analysis was performed by QIIME software package. Results were classified to the phylum, class, order, family and genus levels. To further estimate the function and metabolic pathway of genes, the evolutionary genealogy of genes (EggNOG) database was used via the PICRUS program.

## 3 Results and discussion

### 3.1 EPS increased methane production

The effect of EPS on methane production during anaerobic digestion was investigated, and it was observed that methane production increased with the increase of EPS dosage (Fig. 1). After a period of 28 days, the maximum production rate of 0.93 mmol/d was obtained in the EPS reactor with a dosage of 2 g/L, which was 36.46% higher than that in the control reactor (Fig. 1B). Additionally, in the early stage of anaerobic digestion (1~10 d), the methane production in the EPS reactor was almost the same as that of the control reactor, indicating that the addition of EPS had no significant influence on the methane production (Fig. 1A). This may be because the granular sludge needed time to adapt to the environmental change caused by external substances (EPS) (Lópezmaury et al., 2008). However, in the later stage of digestion (10~28 d), the higher methane production in the EPS reactor could be attributed to promoting DIET between fermenting bacteria and methanogens by electrochemically active substances such as c-Cyts and humus stored in EPS (Xiao et al., 2017). Interestingly, EPS can also be used as carbon source to promote mass transfer by increasing the relative concentration of the substrate, thereby increasing methane production. However, combined with high-throughput sequencing results and culture conditions (sufficient substrate), indicating that EPS may play a role of conductive materials during anaerobic digestion.

**Fig. 1.**
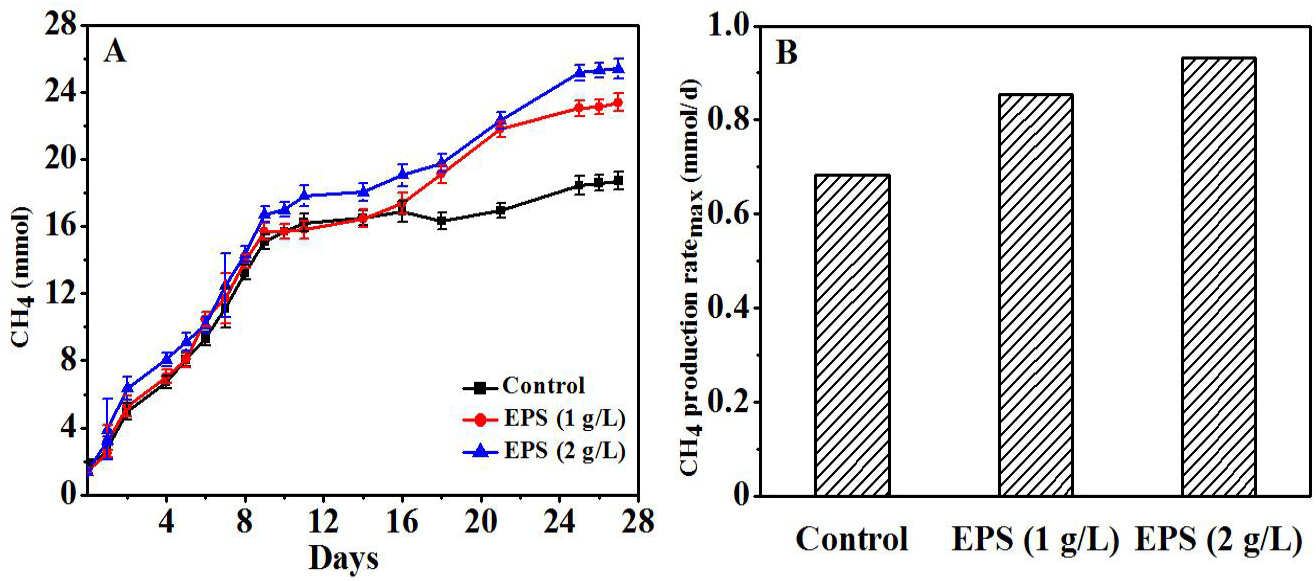
The effect of EPS on the methane production (A); The effect of EPS on the methane production rate_max_ (B).

### 3.2 Effects of EPS on VSS/TSS and particle diameter of AnGS

The ratio of volatile suspended solids (VSS) to total suspended solids (TSS) determined the level of organic components of anaerobic granular sludge (Ding et al., 2015). As shown in Fig. 2A, in the anaerobic digestion process, the VSS/TSS of granular sludge with EPS addition was the same as that of the control reactor, suggesting that the addition of EPS could not affect the composition of the granular sludge. Fig. 2B illustrated that the average particle size of the granular sludge gradually increases as the anaerobic digestion proceeded in the EPS reactor, increasing by 10.31 ± 0.64%. This result was similar to the results reported by Jing *et al.* (2016), in which the mass transfer rate of large granular was more than that of small granular sludge, thus large granular were highly bio-active. Furthermore, the average particle size of granular sludge with EPS addition was approximately equal to that of the control reactor during the anaerobic digestion, illustrating that EPS as an additive could not change the size of granular sludge.

**Fig. 2.**
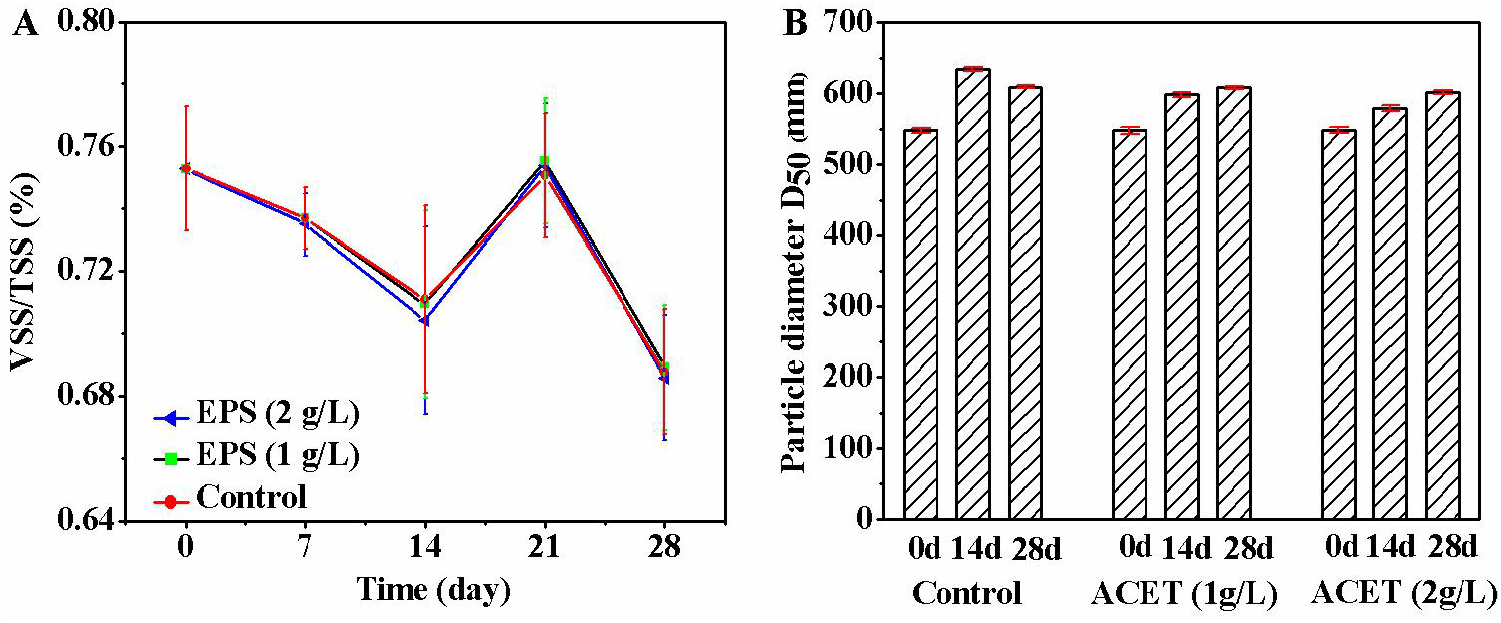
The effect of EPS on VSS/TSS and particle diameter of anaerobic granular sludge.

### 3.3 Effect of EPS as an additive on extracellular polymer substances

Fig. 3A revealed that the concentration of EPS increases as the anaerobic digestion proceeded in all reactors. Moreover, in the EPS reactor, the concentration of extracellular polymer substances was substantially the same as that of the control reactor, indicating that EPS cannot influence the secretion of extracellular polymer substances. The result was different from the conclusion reported by Ye *et al.* (2017), in which the red mud can stimulate microorganisms to secrete a large number of extracellular polymer substances to ensure the normal performance of granular sludge, but the addition of EPS could not cause this phenomenon.

**Fig. 3.**
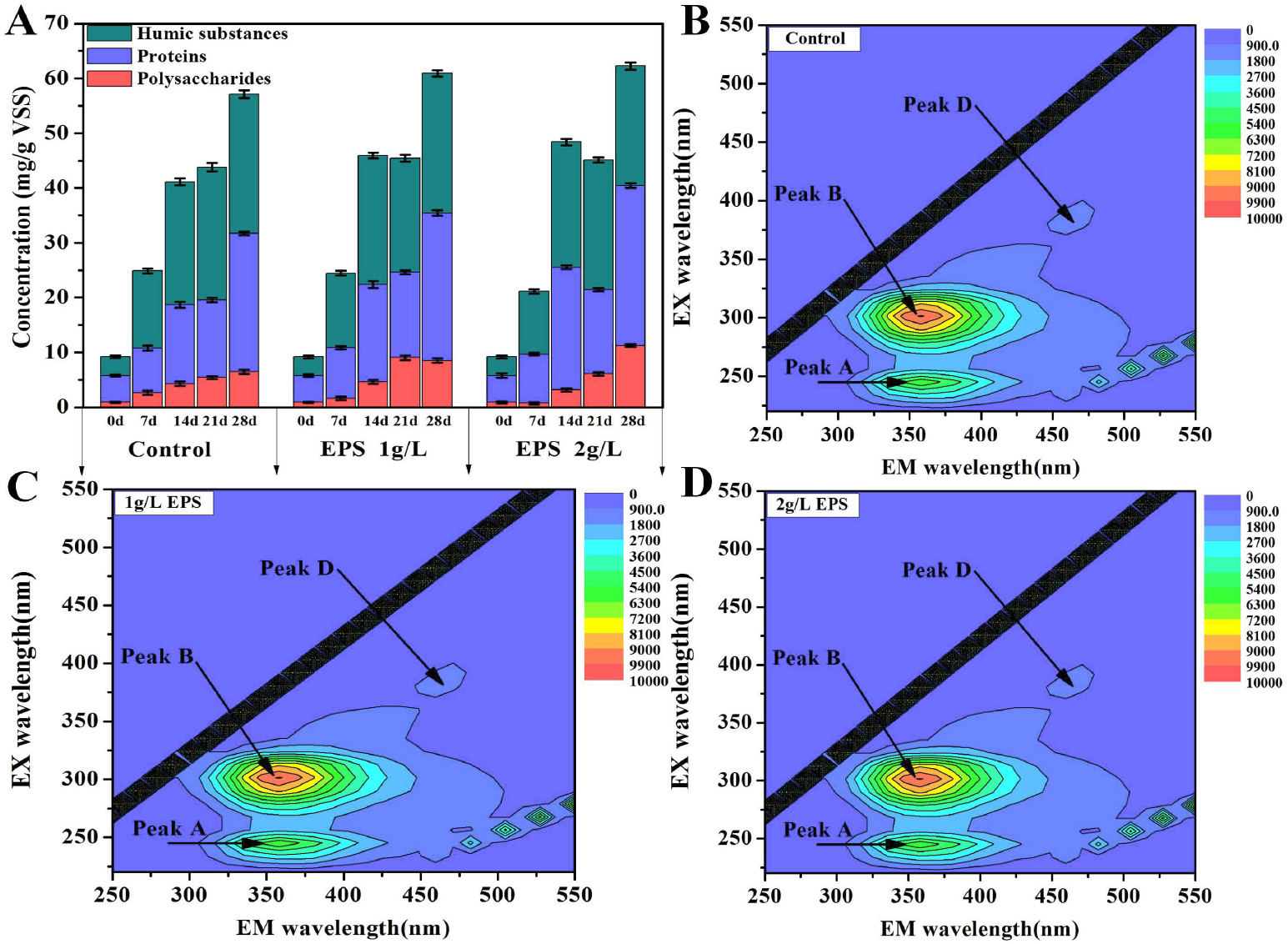
The effect of EPS on the composition and content of extracellular polymer substances.

Fig. 3B-D showed that Peak A, Peak B, and Peak D were the tryptophan-like and soluble microbial by-product-like component (Peak A: Ex/Em=(220, 280)/350) (Yu et al., 2010), protein-like substances (Peak B: Ex/Em=(270, 280)/345-360) (Ouyang et al., 2009), and the humic-like component (Peak D: Ex/Em = (390/(450-470) (Chen et al., 2015; Ding et al., 2015), respectively. Interestingly, the intensities of Peak A, Peak B and Peak D were basically the same in all the reactors, which was in agreement with the conclusions from Fig. 3A.

c-Cyts stored in EPS played an important role in the electron transfer between fermenting bacteria and methanogens (Ye et al., 2018). The absorbance of c-Cyts (419 nm) increased as the anaerobic digestion proceeded in all reactors. More importantly, compared with the control reactor (28 d), the absorbance of c-Cyts was improved by 15.24% in the 2 g/L EPS reactor, suggesting that the addition of EPS can enhance the abundance of c-Cyts (Fig. 4). These results were in step with the conclusion discovered by Ye *et al.* (2017) and Zhang *et al.* (2017), in which the stronger the abundance of c-Cyts, the stronger the ability of transfer electrons. Consequently, we speculated that the higher methane production in the EPS reactor may be due to the increased abundance of c-Cyts.

**Fig. 4.**
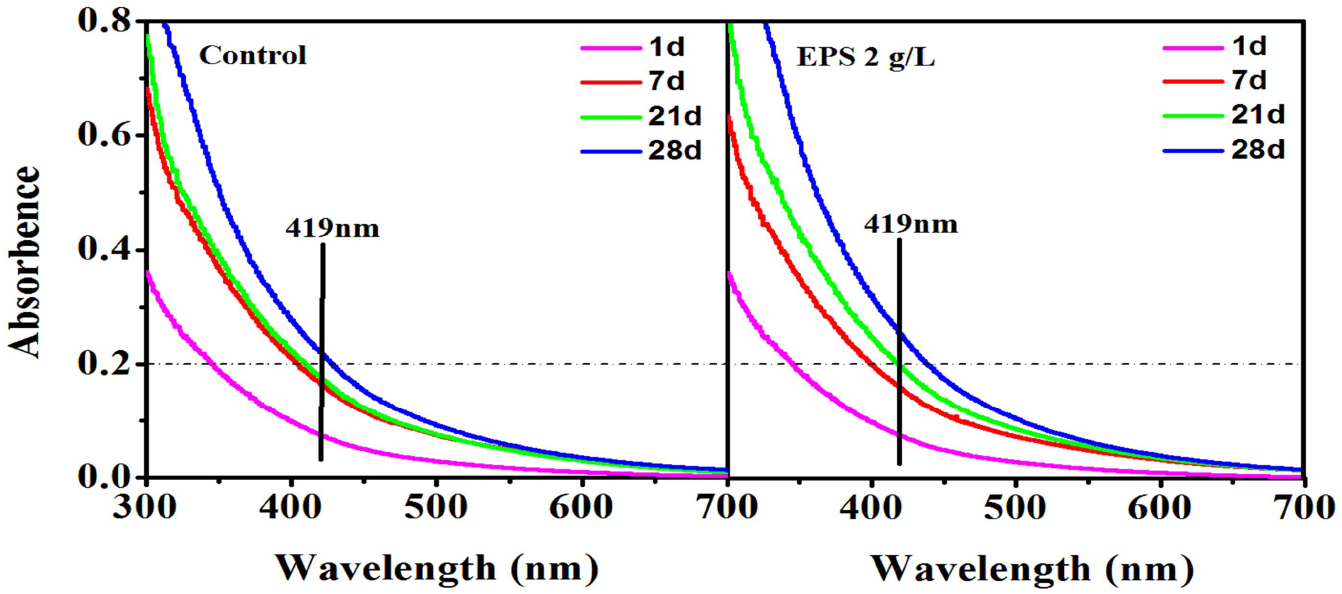
UV/Vis spectra of EPS on days 1, 7, 21, and 28 in the control reactor and the EPS reactor with a dosage of 2 g/L.

### 3.4 Effect of EPS on microbial community structure

#### 3.4.1 Alpha diversity

Table. 1 indicated that the diversity and the number of species in the EPS reactor were higher than that of the control reactor. Taking the Shannon index as an example, compared with the control reactor, the maximum increase was 40.47% in the 2 g/L EPS reactor. Both the diversity and the number of species significantly increased in the EPS reactor because EPS as a potential organic carbon and energy source could change the concentration and type of substrate (Dicker. 2011). In brief, the higher the diversity and the core species in the anaerobic reactor, the better the anaerobic digestion performance of the sludge (Qin *et al.*, 2019). Therefore, the diversity and the number of species significantly increased as a possible factor in the improved of methane production.

**Table 1.**
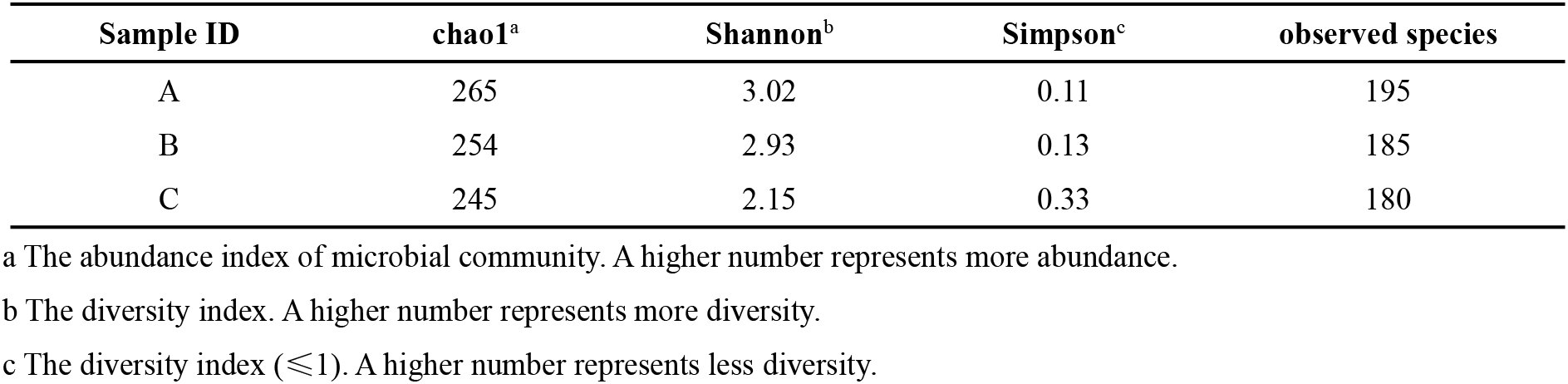
Species diversity and abundance index.

### 3.4.2 Microbial species composition

As shown in Fig. 5, at the phylum level, the microbial composition of granular sludge was basically the same in all reactors, with *Euryarchaeota* as the dominant archaea and *Chloroflexi, Proteobacteria, Actinobacteria, Firmicutes, Synergistetes*, and *Bacteroidetes* as the dominant bacteria. Compared with the control reactor, the relative abundance of *Firmicutes* (38.63%), *Synergistetes* (14.73%) and *Chloroflexi* (6.65%) in the 2 g/L EPS reactor increased by 90.07%, 210.76% and 86.80%, respectively. In addition, the abundance of *Actinobacteria* increased from 0 to 1.37% (Fig. 5A, C). *Actinobacteria, Synergistetes* and *Chloroflexi* as the famous anaerobic hydrolytic and acidifying phyla (Ahlert et al., 2016; Xie et al., 2016), the increase of their relative abundance may accelerate the hydrolysis-acidification process of granular sludge, illustrating that EPS as an additive could enrich functional microorganisms and enhance methane production. This result was similar to that of the effect of conductive materials (granular activated carbon and hematite) on the microbial community structure, both of which can enrich functional microorganisms and change the microbial community structure (Ye et al., 2018; Zhang et al., 2017). Interestingly, the addition of EPS hardly changed the abundance of archaea compared with the control reactor.

**Fig. 5.**
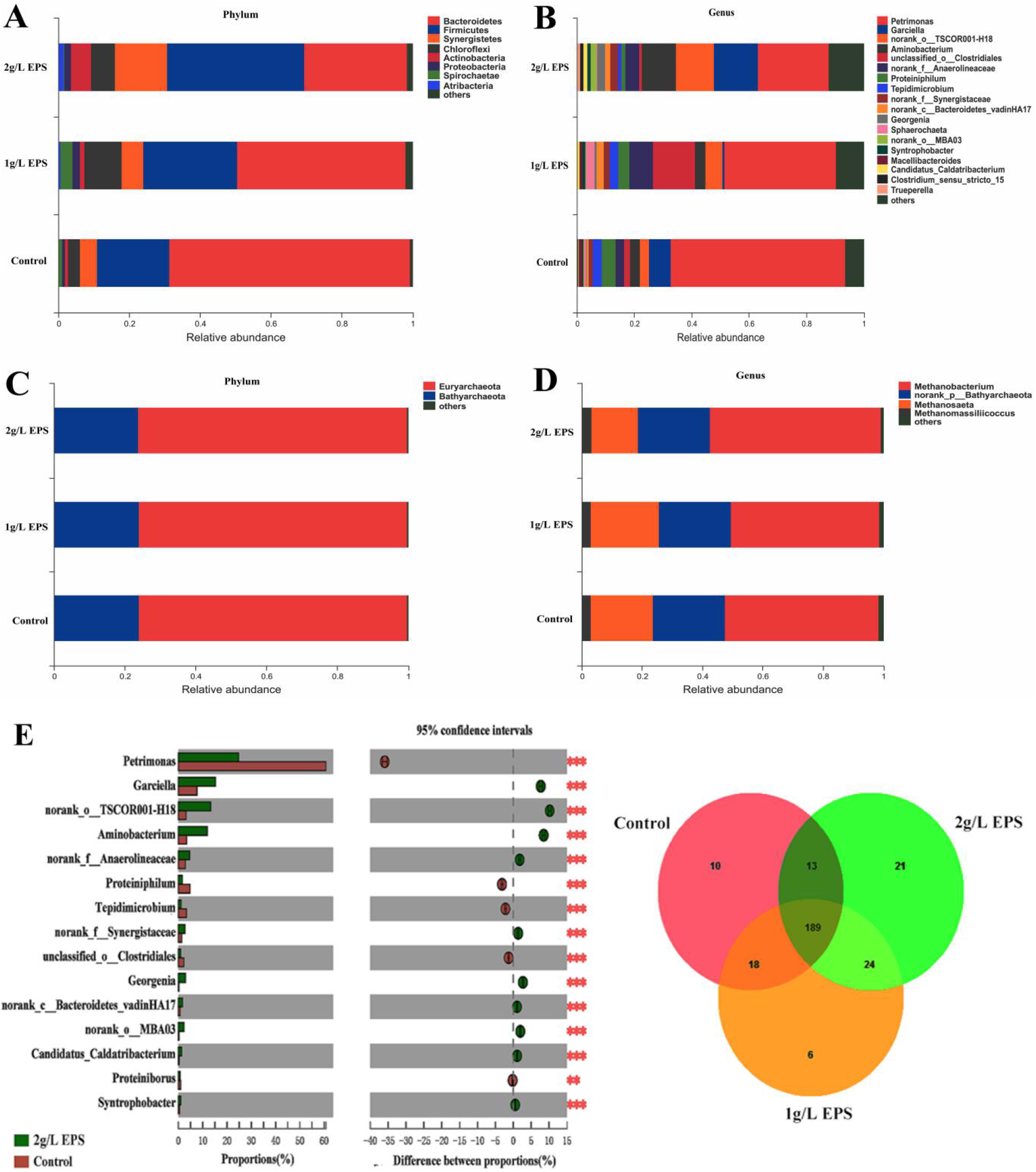
Changes in the predominant microbial community of bacteria and archaea. the distribution of bacteria at the phylum (A) and genus (B) level. the distribution of archaea at the phylum (C) and genus (D) level. Differences between microbial communities in A (2g/L EPS) and C (the control) reactors at the genus level (E).

At the genus level, the bacterial community structure in the EPS reactor changed obviously compared with the control reactor, and the relative abundance of *norank_f__Synergistaceae, Syntrophobacter, Clostridium_sensu_stricto_15* increased significantly (Fig. 5B, D). As for the archaea genus, *Methanobacterium* and *Methanosaeta* were the predominant methanogenic archaea in all reactors. The relative abundance of *Methanobacterium* (methane produced by using H_2_/CO_2_) (Sauer et al., 1980) in the 2 g/L EPS reactor was slightly higher than that in the control reactor. Conversely, the relative abundance of *Methanosaeta* (methane produced by using acetic acid) (Van Haandel et al., 2015) in the 2 g/L EPS reactor was lower than the control (Fig. 5D). This suggested that EPS as an additive could change the methanogenesis pathway of methanogenic archaea. Meanwhile, Fig. 5E also indicated that the addition of EPS can lead to obvious differences in the community structure and composition of bacteria during the anaerobic digestion, which illustrated that EPS mainly affected methanogenesis conducted by changing the microbial community structure and species composition.

To clarify the influence mechanism of EPS on the macrobiotic community structure of granular sludge. The interaction of species was obtained through the correlation analysis of species abundance information of archaea and bacteria in the three reactors, and the formation mechanism of methane yield difference was further explained. Fig. 6 declared that 98.82% OTUs in archaea was related to all anaerobic systems, which verified again the community structure of archaea was basically unchanged. As for the bacteria, 61.97% OTUs was associated with all anaerobic systems; 8.86% OTUs was only related to an anaerobic system with EPS addition, indicating that the macrobiotic community structure of the anaerobic system with EPS addition was significantly different from that of the control system. In addition, 4.92% and 1.64% OTUs were related to an anaerobic system with 2 g/L EPS addition and anaerobic system with 1 g/L EPS addition, respectively. This may be because that the different dosage of EPS in the reactor. More importantly, the correlation between the anaerobic system with EPS addition and control system decreased with the increase of EPS dosage, which was consistent with the conclusions from the Venn diagram. Therefore, we speculated that 8.86% OTUs may be the main reason for the enhanced methanogenic capacity of granular sludge in EPS reactor. Further analysis revealed that 8.86% OTUs from the important hydrolytic and acidification phyla, including *Actinobacteria, Synergistetes* and *Chloroflexi.* Particularly on the carried phase of EPS, its relative abundance was higher than the control, illustrating the possible relationship with EPS carriers. In general, *Synergistetes (norank_f__Synergistaceae, Syntrophobacter, Clostridium_sensu_stricto_15)* and *Methanosaeta* might participate in DIET to enhance methanogenesis via EPS in this study because *norank_f__Synergistaceae, Syntrophobacter, Clostridium_sensu_stricto_15 (Bordeleau et al., 2015) and Methanosaeta* had been shown to transfer electrons between cells (Rotaru et al., 2013; Zhang & Ya-Hai., 2015).

**Fig. 6.**
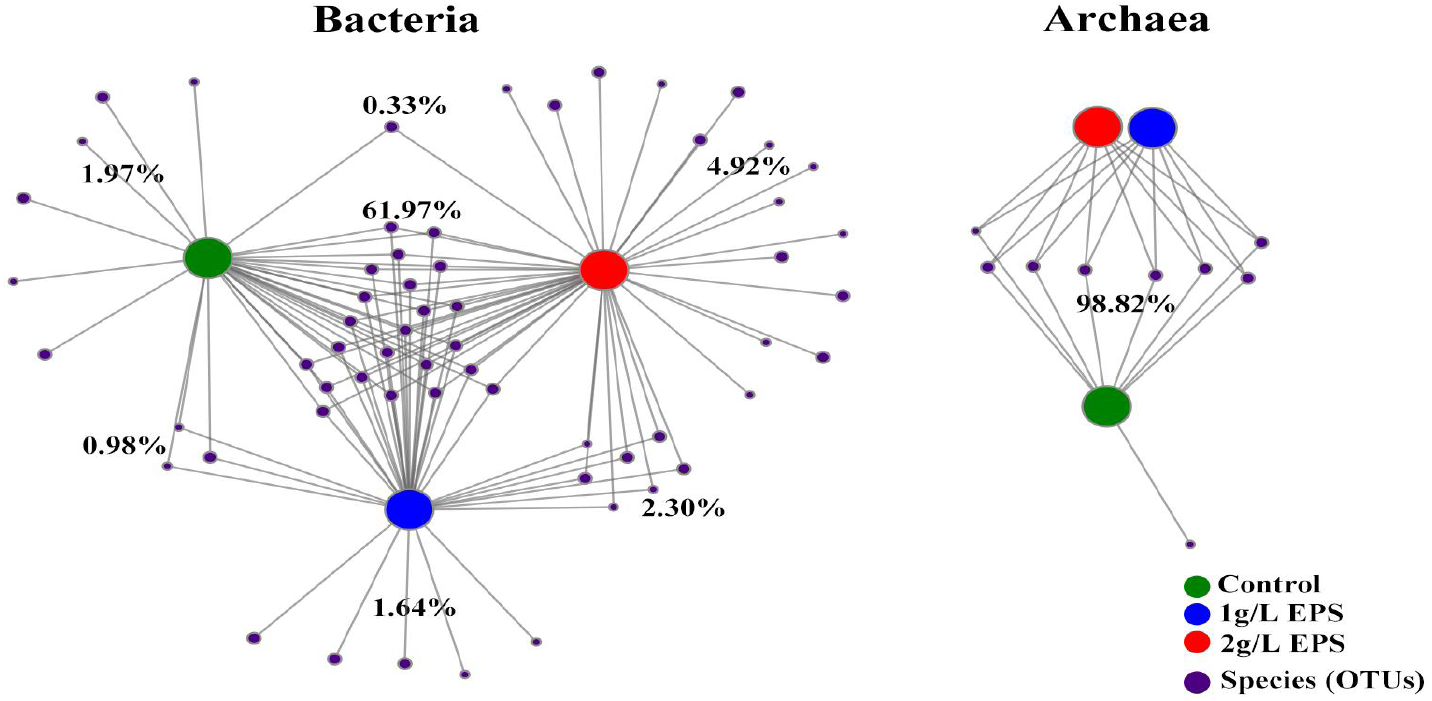
Network analysis of bacteria and archaea on OTU level.

### 3.5 Effect of EPS on methanogenic pathway

The gene samples in the A (2 g/L EPS), B (1 g/L EPS) and C (the control) reactors were compared with the COG database by PICRUSt software, and the results were shown in Fig. 7. The major functional categories in A, B and C reactors included amino acid transport and metabolism (E), general function prediction only (R), cell wall/membrane/envelope biogenesis (M), energy production and conversion (C), and their relative abundances were 8.91, 8.18, 7.85%; 7.97, 8.06, 7.69%; 7.53, 8.08, 9.00%; 7.05, 6.58, 6.12%, respectively. Interestingly, the relative abundance of genes classified in cell wall/membrane/envelope biogenesis (M) was the highest in the control group. The gene abundance of other functions increased with the addition of EPS, which indicated that EPS could change the functional abundance of granular sludge, thereby affected the digestion performance of granular sludge.

**Fig. 7.**
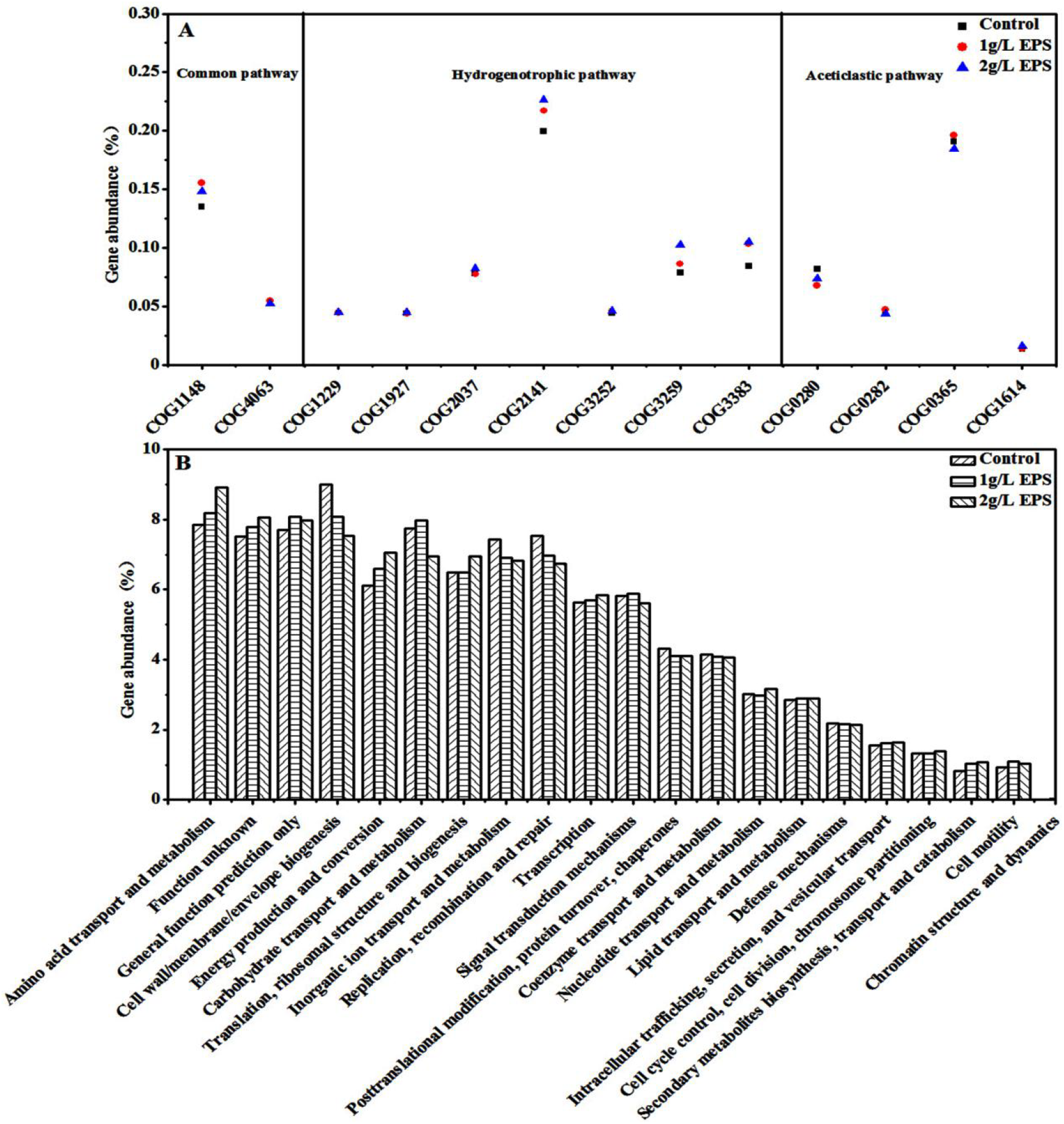
Gene abundance of methane-related enzymes in the three reactors (A). Gene abundance of COG function in the three reactors (B).

The methanogenic pathway of anaerobic granular sludge can be divided into the hydrogenotrophic pathway, acetotrophic pathway, and methylotrophic pathway (Demirel & Scherer., 2008). Fig. 7A showed that the enzymes involved in the acetotrophic pathway in the A, B and C reactors included acetate kinase (COG0282), phosphotransacetylase (COG0280), acetyl-CoA synthetase (COG0365), and their gene abundances were 0.32%, 0.32% and 0.33%, respectively. Interestingly, the gene abundance of enzymes involved in the hydrogenotrophic pathway in the A, B and C reactors was 0.65%, 0.62% and 0.57%, respectively, which indicated that EPS can increase the gene abundance of enzymes related to the hydrogenotrophic pathway (8.77~14.04%), and enhance the hydrogenotrophic pathway of AnGS. Furthermore, the gene abundance of enzymes involved in the common pathway in the A, B and C reactors was 0.20%, 0.21% and 0.19%, respectively. This result was consistent with the analysis of the effect of EPS on the community structure of microorganism.

## 4 Conclusions

This research showed that EPS can increase remarkably methane production in the anaerobic digestion process. Based on the above results, we concluded that the main mechanism of EPS as a biomass conductive material to enhance the methanogenesis of granular sludge was as follows: EPS enriched functional microorganisms such as *Firmicutes, Actinobacteria, Synergistetes, Chloroflexi*, and optimized the community structure of microorganisms. Among them, 8.86% OTUs from the important hydrolysis and acidification phyla, including *Actinobacteria, Synergistetes* and *Chloroflexi*, which may be an important reason for the enhanced methanogenic capacity of AnGS in the EPS reactor. In addition, EPS increased the abundance of c-Cyts, and accelerated the DIET between syntrophic bacteria (*norank_f__Synergistaceae, Syntrophobacter, Clostridium_sensu_stricto_15* increased compared with the control) and methanogens (*Methanosaeta*), thus enhancing the methane production. The 16S function prediction also indicated that EPS could enhance the microbial functional abundance and hydrogenotrophic pathway, improving the anaerobic digestion performance. More importantly, compared with conductive materials (carbon-based materials and conductive minerals), EPS as a biomass conductive material was not only environmentally friendly and economical but also no secondary pollution, low cost and high methane production (Table 2). Therefore, EPS as additives had high research and application value in improving the anaerobic digestion performance.

**Table 2.** Comparison of results with recent studies.

| Additives<sup>a</sup> | Substrate<sup>b</sup> | Digestion mode<sup>c</sup> | Digestion time (day) | Optimal dosage | Biogas yield | Reference |
| --- | --- | --- | --- | --- | --- | --- |
| PAC | CD+WH | SC | - | 4 g/L | ~1.38 L/L of digester per day | (Patel et al., 1992) |
| PAC | CD+PW+CW | SC | - | 4 g/L | ~3.1 L/L of digester per day | (Desai & Madamwar, 1994) |
| Biochar | APL | Batch | 49 | 0.8 g/mL | 5 g/d/L APL | (Torri & Fabbri, 2014) |
| Biochar | sludge | Batch | - | 4 g/L | 16.6 ± 0.6 (mmol CH<sub>4</sub>/g-G) | (Luo et al., 2015) |
| GAC | WAS | Batch | 20 | 5 g/L | increased by 37.2% | (Yang et al., 2017) |
| pine biochar | sludge | sequence | - | 2.49 g/g dry matter of sludge | 0.329 ± 0.001 (mL CH<sub>4</sub>/g COD degraded) | (Shen et al., 2016) |
| Biochar | sludge | SC | 20 | 5 g/L | 127.7 ± 2.7 mL/h | (Zhao et al., 2016) |
| Rud mud | sludge | Batch | 28 | 20 g/L | 1.41 ± 0.02 mmol/g VSS | (Ye et al., 2018) |
| EPS | G | Batch | 28 | 2 g/L | increased by 36.5% | This work |

a PAC: powdered activated charcoal;
b CD: cattle dung; WH: water hyacinth; PW: poultry waste; CW: cheese whey; APL: aqueous pyrolysis liquid; G: glucose;
WAS: wasted activated sludge; c SC: semi-continuous;

## Acknowledgments

The authors are grateful for grants from the National Natural Science Foundation of China (31660182, 21868004), and the project of Guangxi Autonomous Region Department of Science and Technology, China (2017GXNSFAA198200) that supported this research.

